# Effect of luminal surface structure of decellularized aorta on thrombus formation and cell behavior

**DOI:** 10.1101/2021.01.18.427096

**Authors:** Mako Kobayashi, Masako Ohara, Yoshihide Hashimoto, Naoko Nakamura, Toshiya Fujisato, Tsuyoshi Kimura, Akio Kishida

## Abstract

As the number of cardiovascular diseases increases, artificial heart valves and blood vessels have been developed. Although cardiovascular application using decellularized tissue have been studied, the mechanism of their functionality is still unknown. To find the important factor for preparing decellularized cardiovascular prothesis which shows good *in vivo* performance, the effect of luminal surface structure of decellularized aorta on thrombus formation and cell behavior was investigated. Various luminal surface structures of decellularized aorta were prepared by heating, drying and peeling. The luminal surface structure and collagen denaturation was evaluated by immunohistological staining, collagen hybridizing peptide (CHP) staining and scanning electron microscopy (SEM) analysis. To evaluate the effect of luminal surface structure of decellularized aorta on thrombus formation and cell behavior, blood clotting test and recellularization of endothelial cells and smooth muscular cells were performed. The results of blood clotting test showed that the closer the luminal surface structure is to native aorta, the higher the anti-coagulant property. From the result of cell seeding-test, it was suggested that vascular cells recognize the luminal surface structure and regulate adhesion, proliferation and functional expression. These results will provide important factor for preparing decellularized cardiovascular prothesis and lead to future development on decellularized cardiovascular applications.

## 1. Introduction

Cardiovascular disease is currently one of the worldwide leading cause of death(1, 2). Due to a shortage of available donor organs and limitation of the current artificial cardiovascular prothesis, artificial heart valves and vessels with anti-thrombotic, anti-infectious, durable, growth potential and no need for the re-replacement surgery are desired.

Recently, decellularized tissues which are the extracellular matrix obtained by removing cellular components from living tissues have been widely developed. They have gained increasing interest as broad applications as regenerative medicine(3). Although the development of decellularized cardiovascular tissues have been studied(4–6), few of them are clinically applied(3). One of the reasons of this situation is that the mechanisms of the high biocompatibility and functionality of decellularized tissues are not yet understood.

Previously, we reported that decellularized aorta prepared by high-hydrostatic pressurization (HHP) showed good *in vivo* performance, including early reendothelialization and anti-thrombogenicity(7). HHP method disrupts the cells inside the tissue and the cell debris can be removed by series of washing process without using any surfactants. We proposed the hypothesis that vascular endothelial cell recruitment and anti-thrombogenicity occur on the HHP treated aortas since its luminal surface structure is maintained. Therefore, the purpose of this study is to clarify the effect of luminal surface structure on thrombus formation and cell behavior using HHP decellularized aortas. To prepare decellularized aorta with various luminal surface structures, the luminal surface of decellularized aortas were disrupted by heating, drying and peeling. The luminal surface structure was evaluated by immunohistological staining, collagen hybridizing peptide (CHP) staining and scanning electron microscopy (SEM) analysis. To evaluate the effect of luminal surface structure of decellularized aorta on thrombus formation and cell behavior, blood clotting test and recellularization of endothelial cells and smooth muscular cells were performed. These results will provide important factor for preparing decellularized cardiovascular prothesis and lead to future development on decellularized cardiovascular applications.

## 2. Materials and Methods

### 2.1. Preparation of decellularized porcine aorta

Fresh porcine aortas were purchased from a local slaughter house (Tokyo Shibaura Zouki, Tokyo, Japan) and stored at 4 °C until use. The aortas were washed with saline and the surrounding tissue and fat were trimmed. The trimmed aortas were then cut along the longitudinal direction. These aortas were circularly fabricated with an inner diameter of 15 mm with a hollow punch. The disked aortas were packed in plastic bags with saline and hermetically sealed. After the cells were destroyed by hydrostatically pressurization at 1000 MPa and 30 °C for 10 min using a hydrostatic pressurization system (Dr. Chef, Kobelco, Tokyo, Japan), samples were then washed with DNase (0.2 mg/mL) and MgCl_2_ (50 mM) in saline at 4 °C for 7 d, followed by changing the washing solution to 80% ethanol in saline at 4 °C for 3 d, and with saline at 4 °C for 3 d to remove cell debris in the tissues.

To prepare decellularized aorta with various luminal surface structures, some HHP decellularized aortas were placed in the sterile flask with saline and put in the heating mantle at 90 °C for an hour (hereinafter called HHP 90 °C tunica-intima). Some of the aortas were placed on the sterile drape with the inner surface up and dried for 30 min (hereinafter called HHP 30 min dried tunica-intima). For some decellularized aortas, the inner membrane were peeled off to expose the tunica-media (hereinafter called HHP tunica-media).

### 2.2. DNA quantification

The decellularized aortas were freeze-dried and dissolved in a lysis buffer containing 50 μg/mL protease K, 50 mM Tris-HCl, 1% SDS, 10 mM NaCl, and 20 mM EDTA at 55 °C for overnight. DNA extraction and purification was performed with phenol/chloroform and ethanol precipitation. The residual DNA content in the native tissue and decellularized tissue were quantified using Quant-iT PicoGreen ds DNA reagent (Thermo Fisher Scientific K. K., Tokyo, Japan) against a λDNA standard curve (0–1000 ng/mL, Thermo Fisher Scientific K. K., Tokyo, Japan) using a microplate reader (excitation 480 nm, emission 525 nm, Cytation 5, BioTek Instruments, Inc., Vinuski, VT, USA). The measurements were normalized to the tissue dry weight of 20 mg.

### 2.3. Histological evaluation of decellularized aortas

The native aorta and decellularized aortas were fixed by immersion in a neutral-buffered (pH 7.4) solution of 10% formalin in PBS for 24 h at room temperature and dehydrated in graded ethanol. The samples were then immersed in xylene and embedded in paraffin. Paraffin sections were cut into 4 μm-thick sections for H-E and and 5 μm-thick sections for anti-type IV collagen staining.

### 2.4. F-CHP staining

After paraffin was removed by rinsing with xylene and graded ethanol, slides were rinsed with deionized water twice and with 1 x PBS thrice to remove detergents. A solution containing 15 μm of F-CHP diluted with PBS was heated at 80 °C for 5 min to dissociate trimeric CHP to monomers. The heated CHP solution was cooled immediately in an ice bath to avoid thermal damage to the tissue sections. The CHP solution was dropped onto each section and the samples were incubated in a humidity chamber at 4 °C for 2 h. The slides were then washed three times with PBS before mounting.

### 2.5. Scanning electron microscope (SEM) observation

A scanning electron microscope (S-4500/EMAX-700, Hitachi, Ltd.) was used. The aortas were fixed with 2.5% glutaraldehyde in PBS and dehydrated gradually in ethanol. Dehydrated samples were placed in tert-butyl alcohol and then vacuum dried. Before observation, the surfaces of the decellularized aortas were coated with gold.

### 2.6. Evaluation of thrombogenicity of decellularized aortas

Blood coagulation test was performed following a previously described protocol(8). Briefly, whole blood containing 0.324% citric acid was prepared with a tenth of calcium chloride (CaCl_2_) to coagulate blood for 15 min on glass. A stainless-steel tray covered with a moistened paper towel was floated in a 37 °C water bath. Whole blood was dropped onto cover glasses that were lined up on the tray. Each cover glass was picked up at 2, 4, 6, 8, 10, and 15 min and washed with saline, and then checked for whether the whole blood was coagulated or not at 15 min. When the concentration of CaCl_2_ was decided, the same amounts of CaCl_2_ and whole blood were mixed. In the same way, decellularized aortas were placed on the tray and 50 µL of whole blood containing CaCl_2_ was added. These samples were washed with saline at the same intervals of time described above and photographs were taken. All decellularized aortas were then placed into a 24-well plate with deionized water, resulting in hemolysis. The absorbance of the hemolyzed blood was measured with an absorbance meter 24 h later.

### 2.7. Cell seeding

The disked aortas were placed on a 24-well tissue culture plate and a stainless-steel ring with a 13 mm inner and 15 mm outer diameter was placed onto them to avoid the samples from curling. Human umbilical vein endothelial cells (HUVECs) and human aortic smooth muscle cells (HAoSMCs) were purchased from TAKARA BIO (Tokyo, Japan). HUVECs were seeded at 2 × 10^4^ cells/cm^2^ and HAoSMCs were seeded at 1 × 10^4^ cells/cm^2^ on the surface of the decellularized aortas and incubated at 37 °C under 5% CO_2_ conditions.

### 2.8. Measurement of cell proliferation

HUVECs and HAoSMCs were stained with calcein-AM and then incubated for 30 min at 37 °C. The cells were observed using a fluorescent microscope (BZ-X710, Keyence Corp., Osaka, Japan). The numbers of cells were counted in sections from all samples and the cell density was calculated using the counted cell numbers.

### 2.9. Quantitative reverse-transcription Polymerase Chain Reaction (qPCR)

Total RNA was extracted from HAoSMCs on TCPS and decellularized aortas using according to the manufacturer’s instructions. GAPDH was used to normalize gene expressions. Quantitative PCR was performed using Power SYBR Green PCR Master Mix (Thermo Fisher Scientific) on a StepOnePlus system (Thermo Fisher Scientific) with Delta Delta Ct method. Forward and reverse primer sequences are shown in table.

### 2.10. Statistical Analysis

The quantitative analysis of residual DNA and cell density are expressed as the mean ± standard deviation (SD) and the blood coagulation rate data is expressed as the mean ± standard error of the mean (SE). Analysis of variance followed by Student’s *t*-test was used to determine significant differences among groups.

## 3. Results and Discussion

Porcine aortas were decellularized and the amount of residual DNA was measured (Fig 1). The amount of DNA remaining in the HHP treated aorta was significantly lower than untreated aorta.

**Figure 1.**
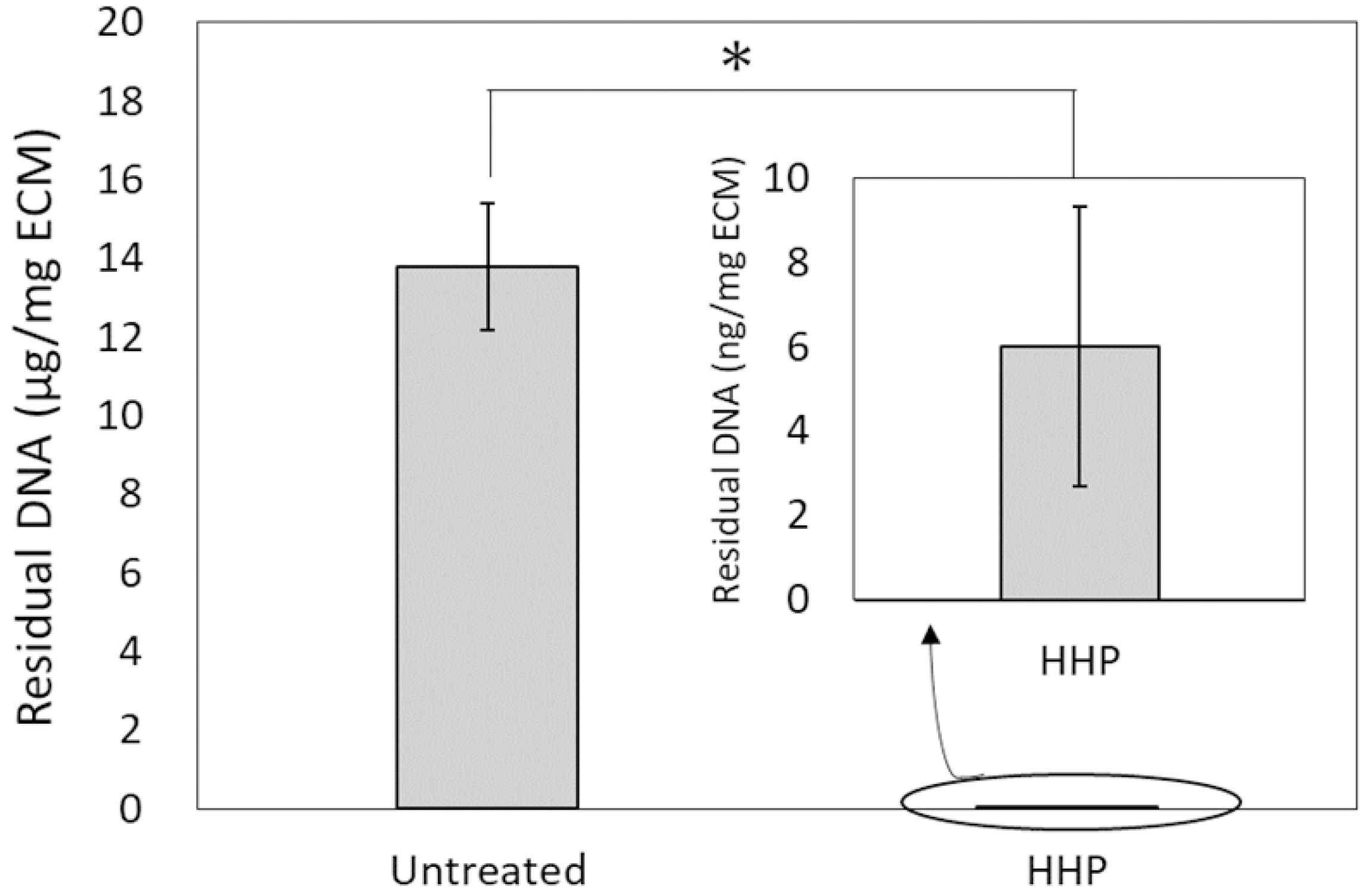
Quantitative analysis of residual double stranded DNA (dsDNA) in decellularized aortas. \**p* < o.oo1.

Figure 2 (A) - (J) showed H-E and Type IV collagen-stained native aorta and decellularized aortas. The efficacy of decellularization were verified by H-E staining. No nuclei were detected in sections of decellularized aortas by HHP treatment (Figure 2 (B) - (E)). The tunica-intima of the HHP treated aorta and the HHP 30 min dried treated tunica-intima was observed similarly to that of the untreated aorta (Fig. 2 (B), (D)). The 90 °C heated tunica-intima of HHP treated aorta had larger wave-like shapes and a flatter fibril structure than native aorta(Fig.2(C)). In the tunica-media of HHP treated aorta, the gaps between collagen fibrils were enlarged due to the peeling of tunica-intima (Fig.2(E)). The immunostaining of type IV collagen was also performed. The type IV collagen is the main component of the basement membrane and forms networks that provide the structural support for endothelial cells. The type IV collagen layer was preserved at HHP decellularized tunica-intima, even they were dried (Fig. 2 (G),(I)). HHP 90 °C tunica-intima was weakly stained (Fig.2(H)).

**Figure 2.**
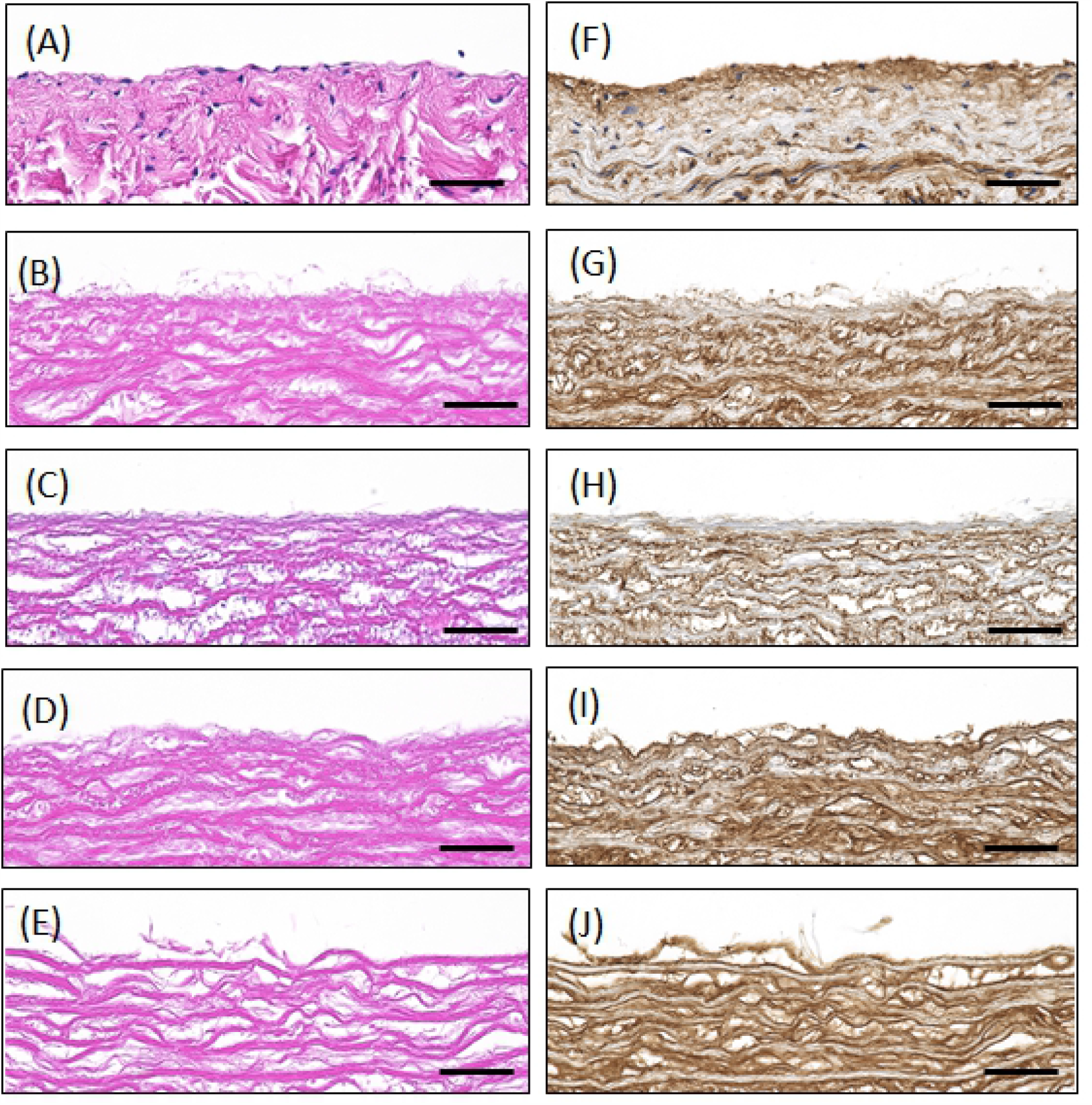
H-E staining (A-E), type IV collagen staining(F-J) of decellularized porcine aortas. (A)(F)Untreated aorta, (B)(G)HHP tunica-intima, (C)(H)HHP 90 °C tunica-intima, (D)(I) HHP 30min dried tunica-intima, (E)(J) HHP tunica-media.

**Figure 3.**
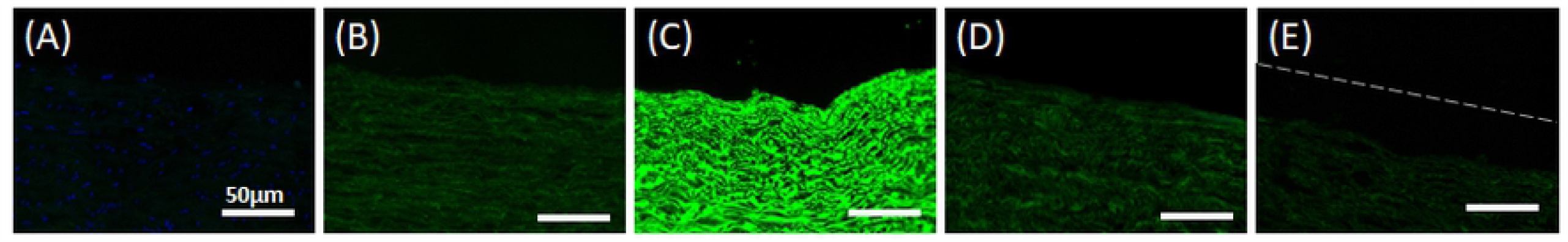
Fluorescence images showing CF-CHP staining on sections of (A) Untreated aorta, (B) HHP tunica-intima, (C) HHP 90 °C tunica-intima, (D) HHP 30 min dried tunica-intima, (E)(J) HHP tunica-media. Scale bar : 50μm.

**Figure 4.**
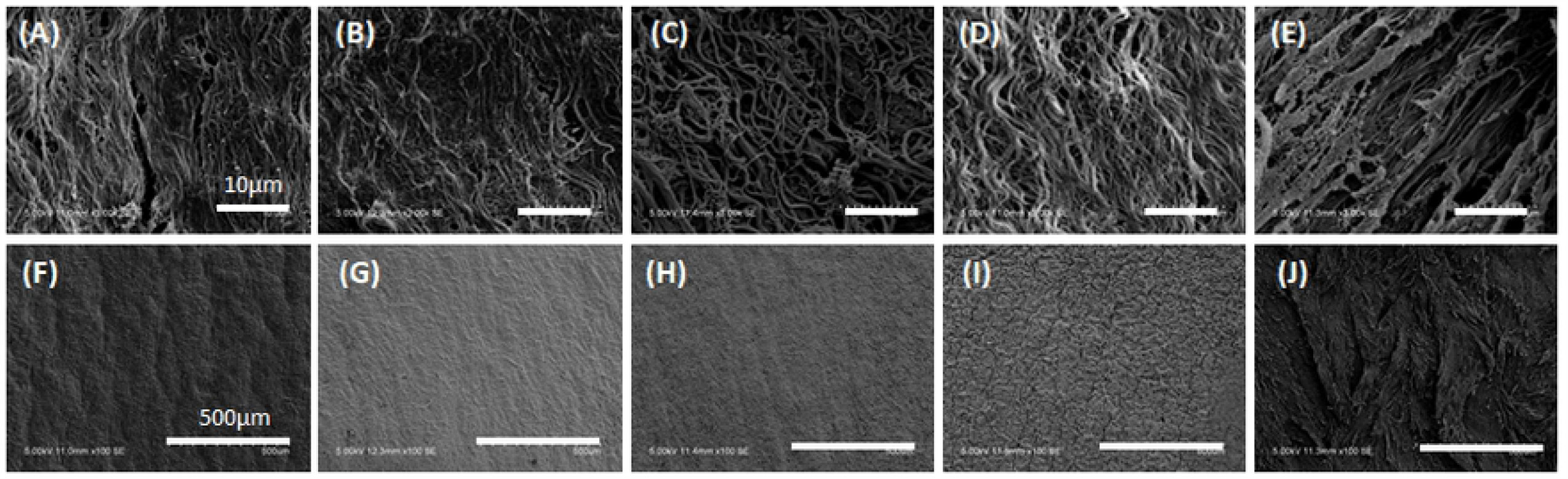
SEM observation of surface. (A)(F) Untreated aorta, (B)(G) HHP tunica-intima, (C)(H) HHP 90 °C tunica-intima, (D)(I) HHP 30 min dried tunica-intima, (E)(J) HHP tunica-media. Scale bar: (A)-(E)10 μm, (F)-(J) 500 μm.

The Collagen Hybridizing Peptide (CHP) staining which specifically target denatured collagen chains was also performed. As expected, no fluorescence intensity except DAPI was detected from untreated aorta (Fig.3 (A)). HHP treated tunica-intima showed almost no intensity-signal, suggesting slightly denatured collagen (Fig.3(B)). In HHP 90 °C tunica-intima, strong fluorescence signal was detected, indicating that the collagen was completely destroyed due to the heating process (Fig.3(C)). No significant difference was observed between the fluorescence image of HHP tunica-intima and that of HHP 30 min dried tunica-intima and HHP tunica-media (Fig.3(D),(E)).

SEM was used to analyze the fiber structure of decellularized aortas (Fig.4). The fibers were observed in magnified 3.0k SEM images (Fig.4(A)-(E)), while whole surface image was observed in 100 SEM images (Fig.4(F)-(J)). For HHP treated tunica-intima, the fiber bundles were oriented longitudinally and similar to the untreated aortas (Fig.4 (B), (G)). As for HHP 90 °C tunica-intima, thick and shrunken fibers were observed in the magnified images (Fig.4 (C)) and smooth plane surface without ruggedness were shown in Fig.4(H). Compared to the SEM images of HHP tunica-intima, gaps between fibers (Fig.4(D)) and a surface with fine cracks were observed in the HHP 30 min dried tunica-intima (Fig.4(I)). The HHP treated tunica-media showed rough surface due to the peeling of tunica-intima (Fig.4 (E), (J)). From above observation, various luminal surface structure of decellularized aortas were prepared.

In this study, Lee-White test was used to examine the effect of luminal surface structure on the thrombogenicity of decellularized aortas. Figure 5 (A) exhibits picture of *in vitro* evaluation test of blood clotting test for decellularized aortas. Glass and PTFE were used as negative and positive controls, respectively. On glass, the blood clot formed at 4 min, while no clot formation occurred until 15 min on PTFE. As for HHP treated tunica-intima, no clot formation was observed until 15 min, which is the almost same result as PTFE. As for HHP 90 °C tunica-intima, HHP 30 min dried tunica-intima and HHP treated tunica-media, the thrombus formation occurred within 10 min. The result of Figure B shows the measurement of absorbance of those hemolyzed blood clot. As shown in Figure.5(B), HHP 90 °C treated tunica-intima, HHP 30 min dried tunica-intima and HHP treated tunica-media showed nearly the same result. The thrombogenicity of HHP decellularized aorta was comparable with PTFE.

**Figure 5.**
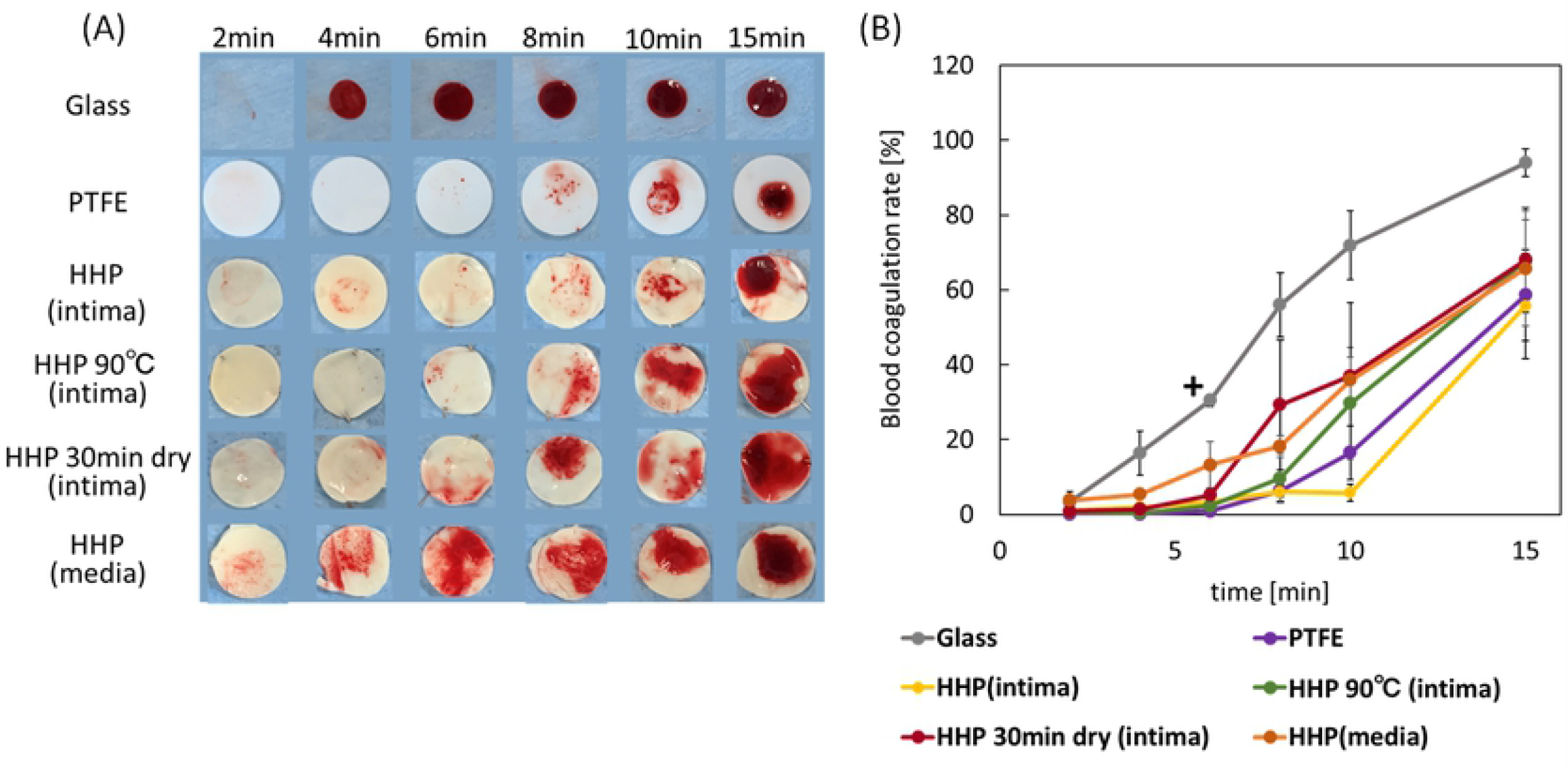
(A) Photograph of whole blood coagulation time of decellularized aortas with various basement membrane structure. (B) Blood coagulation rate of decellularized aortas. Error bars represent S.E. The cross (+) shows a significant difference (p<0.05) between glass and PTFE, HHP (intima), HHP90 °C (intima), HHP 30min dry (intima) at 6 min.

It is widely known that endothelial cell seeding on luminal surface of cardiovascular grafts are required to prevent thrombus formation(9, 10).

Endothelial cells also regulate vessel tone, platelet and leukocyte activation and SMCs migration and proliferation(11). In this study, HUVECs were chosen to evaluate the recellularization efficacy and cell behavior on decellularized aortas with various luminal surface structure. The attachment and proliferation of HUVECs on decellularized aortas were observed by fluorescence microscope. The numbers of cells were then counted in sections from all samples and the cell density was calculated using the counted cell numbers (Fig.6(K)). HUVECs adhered on all samples. HUVECs were well-proliferated on HHP treated tunica-intima (Fig.6 (B), (G), (K)). The attachment of HUVEC on HHP 90 °C tunica-intima and HHP 30min dried tunica-intima were nearly equal to that of the HHP treated tunica-intima, but cells did not proliferate and extend (Fig 6 (C),(D),(H),(I),(K)). On HHP treated tunica-media, initial adhesion of HUVEC was low and the number of HUVECs decreased during 7 day cultivation (Fig.6 (E),(J), (K)).

**Figure 6.**
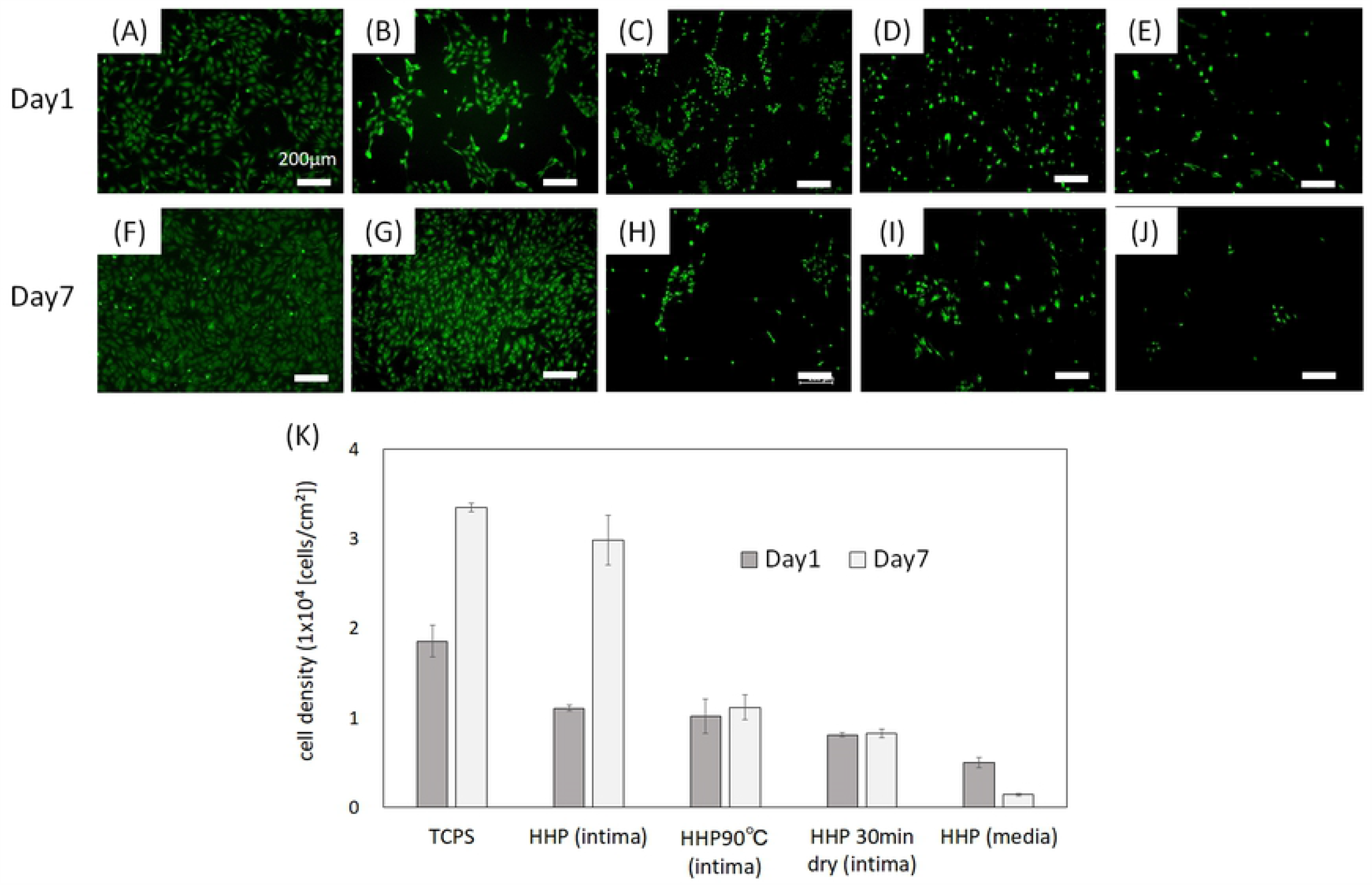
Fluorescence images of HUVEC at day1 and day7 on (A)(F)TCPS, (B)(G) HHP decellularized tunica-intima, (C)(H)HHP decellularized 90 °C heated tunica-intima, (D)(I) HHP 30min dried tunica-intima, (E)(J)HHP tunica-media. (K) Cell density. Scale bar : 200 μm.

Vascular smooth muscle cells (VSMCs) are present in the tunica media (middle layer) and have secretory ability of the major extracellular proteins including collagen, elastin and proteoglycans that involves mechanical properties(12).Furthermore, VSMCs can switch between contractile phenotype and synthetic phenotype in response to changes in local environment(13, 14)*

They usually exhibit contractile phenotype and express contractile proteins such as *α*-smooth muscle actin (SMA), smooth muscle calponin (CNN) and SM22 *α* (SM22) and calponin. In contractile phenotype, they have a low proliferative rate to maintain the ECM of tunica-media(13). In response to vascular injury including tissue damage, they alter their phenotype to synthetic state and reduce the expression of contractile proteins, with increasing proliferation and remodeling the ECM to facilitate migration(15–17). In this study, the effect of the luminal surface structure of decellularized aorta on VSMC’s proliferation and phenotype were investigated. Human aorta smooth muscle cells (HAoSMCs) were seeded on TCPS and decellularized aortas. The cell density was calculated by counting cell numbers on each sample (Fig.7 (K)). From Fig 7(A) to (J), HAoSMCs adhered on all samples. HAoSMCs were well-proliferated on HHP treated tunica-intima and HHP 30min dried tunica-intima (Fig.7 (B), (D), (G), (I),(K)). The initial attachment of HAoSMCs on HHP 90 °C tunica-intima and HHP tunica-media were nearly equal to that of the HHP treated tunica-intima, but cells hardly proliferated (Fig.7 (C), (E), (H), (J),(K)).

**Figure 7.**
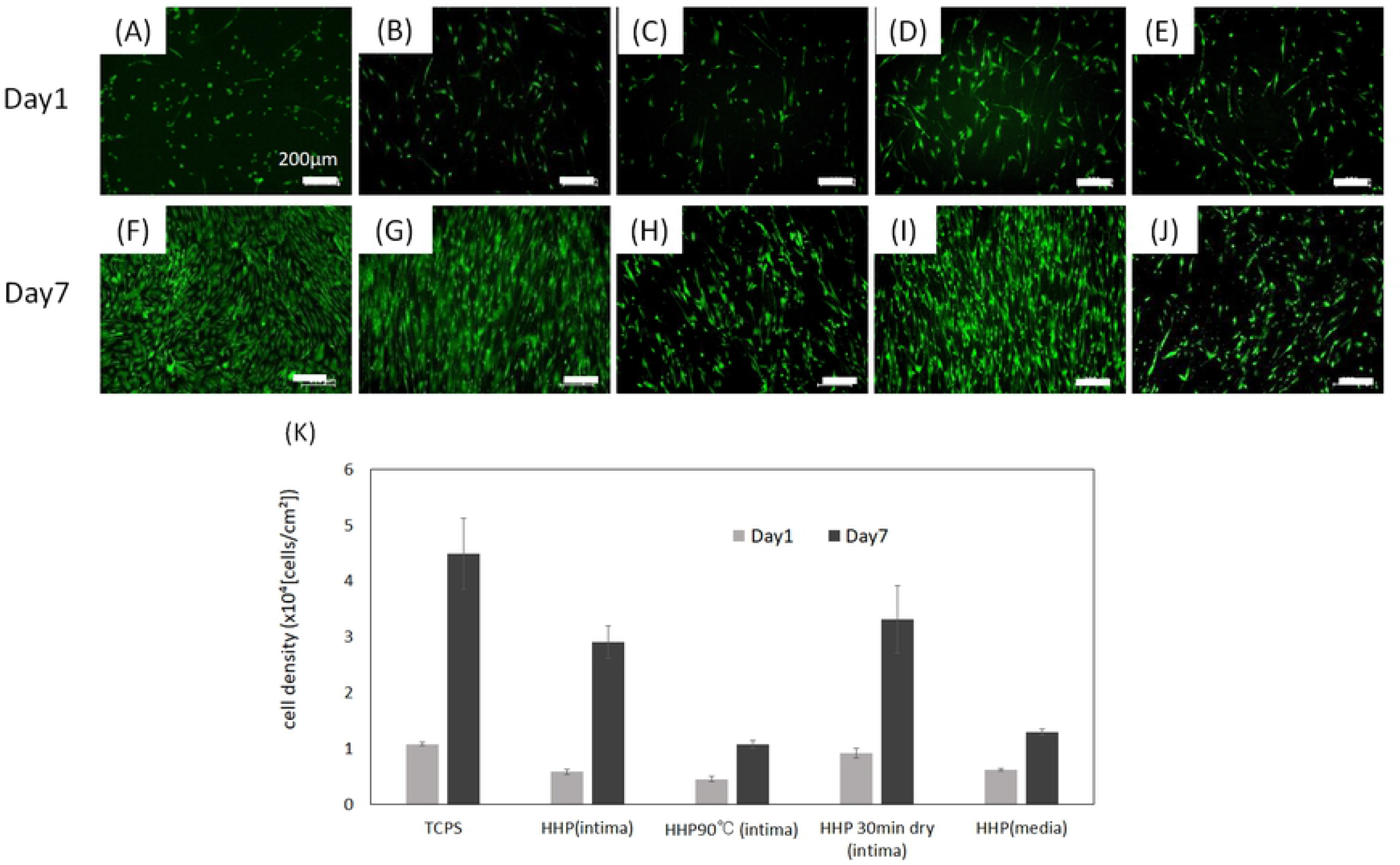
Fluorescence images of HAoSMC at day1 and day7 on (A)(F)TCPS, (B)(G) HHP decellularized tunica-intima, (C)(H)HHP 90 °C heated tunica-intima, (D)(I)HHP 30min dried tunica-intima, (E)(J) HHP decellularized tunica-media. Scale bar : 200μm.

The level of expression of SMA, CNN and SM22 of HAoSMCs on decellularized aortas was examined by qRT-PCR analysis. On day1, HAoSMCs on all samples showed low gene expression due to a small number of cells (Fig. 8 (A)). On the other hand, on day7, the expression ratio of contractile phenotype genes was high, except at HHP 90 °C heated tunica-intima. Since there was few cell adhesion on HHP 90 °C heated tunica-intima, the amount of gene expression was low (Fig. 8 (B)).

**Figure 8.**
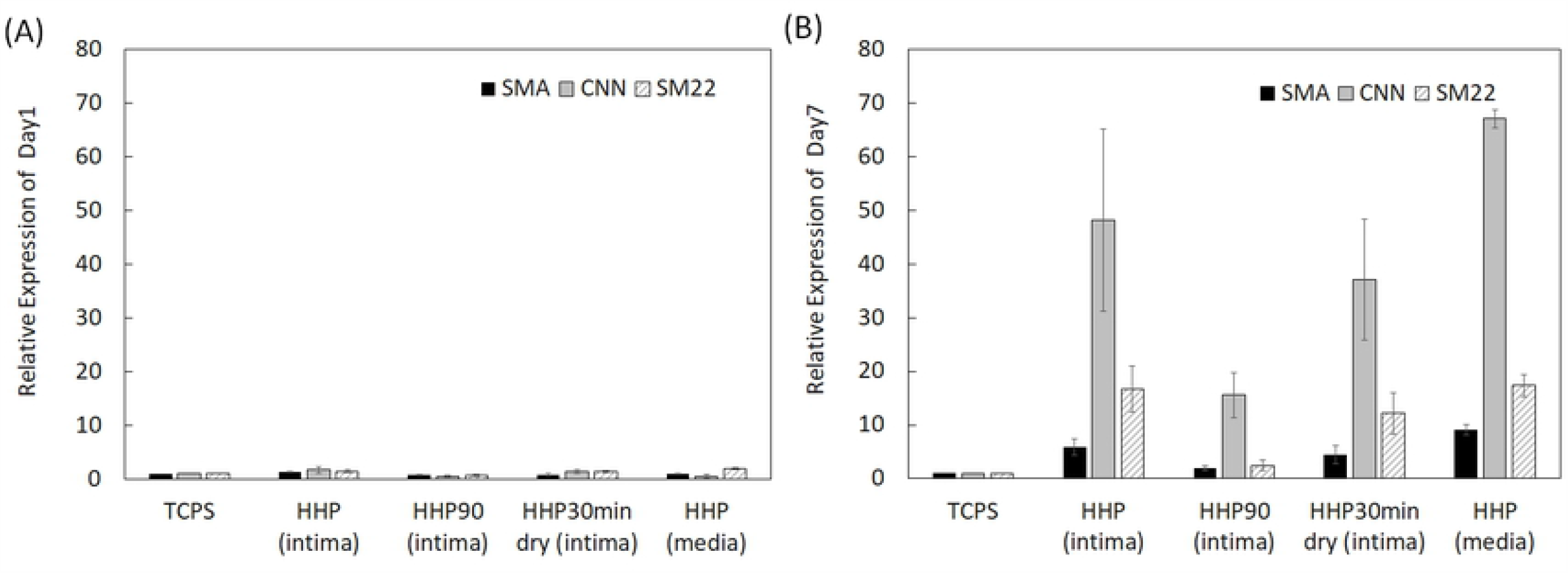
mRNA expression of SMA, CNN and SM22 in HAoSMCs on TCPS and decellularized aortas, (A) at day1 and (B) at day7.

## 4. Discussion

To identify the effect of luminal surface structure of decellularized aorta on thrombus formation and cell behavior, porcine aortas were decellularized using HHP method that have been previously reported good *in vivo* performance. Their luminal surfaces were then fabricated by heating, drying and peeling. The luminal surface structure and components were evaluated by type IV collagen immunostaining, CHP staining and SEM analysis. Type IV collagen immunostaining was performed to evaluate the maintenance of the basement membrane of decellularized aorta. There was no effect of HHP decellularization on type IV collagen, but type IV collagen was not stained after heating process. This result indicates that type IV collagen was denatured and gelatinized by 90°C heating process. CHP staining is known as a method to examine the unfolding of collagen molecules in decellularized tissue(18). Compared to the native aorta, slightly unfolded collagen was observed when aortas were decellularized by HHP method. The image of HHP 90°C heated tunica-intima clearly showed that collagen molecules were completely denatured when they were heated. SEM observation was performed to observe the fiber structure of decellularized aorta. In the HHP 90°C heated tunica-intima, the fibers appeared thicker and shrunken because the collagen fiber was denatured by heat treatment. This result correlates with previous report which showed that the collagen fibers shrink and their fiber diameter increases when they treated with temperatures at or above 50 °C (19, 20). On the other hand, from the SEM images of HHP 30 min dried tunica-intima, drying caused cracks on the tissue surface and the gaps between fibers were clearly visible. From these results, HHP decellularized aortas with different luminal surface structure and collagen denaturation were prepared.

The effect of luminal surface of decellularized aorta on thrombus formation was performed by blood clotting test. From the results, the anti-coagulability was higher for HHP treated tunica-intima and lower for the decellularized aortas with disordered luminal surface structure (HHP 90 °C heated tunica-intima, HHP 30min dried tunica-intima, HHP tunica-media). This indicated that the closer the luminal surface structure is to native aorta, the higher the anti-coagulant property.

The effect of luminal surface of decellularized aorta on vascular cell behavior was performed using HUVECs and HAoSMCs. Endothelial cells and smooth muscle cells are the main cellular components of aortas and interact each other to maintain the function and mechanical property of vessel(21). It is important to evaluate whether those vascular cells which play essential role *in vivo* recognize the place where they should be exist and proliferate and express their functions properly. Cell seeding on decellularized aortas and the evaluation of mRNA expression of those cells by RT-PCR were performed. When HUVECs and HAoSMCs were seeded on HHP decellularized tunica-intima which is similar to the native aorta in both luminal surface structure and components, they adhered and well-proliferated. Since luminal surface structure and basement membrane were well-maintained on HHP decellularized tunica-intima. On the other hand, HAoSMCs showed high proliferation since it is suggested that they recognize the luminal surface as tunica-intima and exhibit synthetic type which shows high proliferative capacity. On the HHP tunica-media, initial adhesion of HUVECs were observed but they sloughed off during cell culture. It may be because the basement membrane which support endothelial cell was removed. HAoSMCs on HHP tunica-media did not proliferated. This result is similar to the vascular smooth cell behavior *in vivo* which has been reported that smooth muscle cells exhibit stable contractile type with low proliferative capacity. Therefore, HAoSMCs recognized the luminal surface structure of tunica-media and exhibit stable contractile type with low proliferative capacity. As for the HHP 90 °C heated tunica-intima which luminal surface structure and collagen were denatured, initial adhesion of both HUVECs and HAoSMCs are low and did not proliferate. It may be because luminal surface structure is disordered and type IV collagen is denatured. As for the HHP 30min dried tunica-intima which luminal surface structure was cracked by drying, HUVECs did not proliferate while HAoSMCs adhered and proliferated. From this result, it was suggested that drying out luminal surface of decellularized aorta may lead to intima hyperplasia. Intima hyperplasia is the thickening of the tunica-intima due to the growth of vascular smooth muscle cells(22). This *in vitro* result is correlated with previous *in vivo* study which shows vessel occlusion when intima were dried before implantation(23). From these results, it is suggested that HUVEC and HAoSMCs recognize the luminal surface of decellularized aorta and proliferate and express their functions properly. Furthermore, the maintenance of luminal surface structure of decellularized aorta is one of the key factors for preparing the decellularized cardiovascular application.

## 5. Conclusion

In the present study, decellularized aorta with different luminal surface structure were prepared to investigate the effect of their structures on thrombus formation and cell behavior. The results of blood clotting test showed that the closer the luminal surface structure is to native aorta, the higher the anti-coagulant property. From the results of cell seeding and qRT-PCR, it was found that endothelial cells and smooth muscle cells recognize where they were seeded and regulate adhesion, proliferation and functional expression. These results provide useful insights into future development on decellularized cardiovascular applications.

## Acknowledgements

This work was supported in part by a Grant-Aid for Scientific Research (B) (16H03180) from JSPS, the Creative Scientific Research of the Viable Material via Integration of Biology and Engineering from MEXT and the Cooperative Research Project of Research Center for Biomedical Engineering from MEXT.

